# Altered *Arabidopsis thaliana* sugar metabolism affects exudation, immune responses, and plant-microbe interactions

**DOI:** 10.1101/2024.09.07.611788

**Authors:** Alexandra Siffert, Joëlle Sasse

## Abstract

Sugars are critical for plant growth, development, and environmental interactions. They have multiple roles as nutrients for plants, associated beneficial and pathogenic microbes, and as signaling compounds for immunity. We characterize the interconnectedness of these functions by analyzing sugar metabolism and transporter mutant lines. We find that in these lines, root-derived compounds, exudates, are significantly altered in comparison with wild-type not only for carbohydrates, but also for lipids, organic acids, and defense compounds. Quantification of sugar exudation reveals more carbon release during the day than at night, altered sugar exudation in mutant lines, and an opposite exudation pattern with elevated exudation at night for *pgm1*, a line deficient in starch synthesis. Sugar levels in exudates and tissues did not correlate, suggesting a controlled mode of exudation for sugars. Altered sugar levels have functional consequences: mutant lines exhibit increased resistance against the pathogen *Pseudomonas syringae* and harbor altered numbers of microbes on roots. Day- and nighttime exudates of mutant lines impact the growth of single microbes such as an inability to grow for *Bacillus subtilis*. Exogenous sugar alters the production of reactive oxygen species in a plant development-dependent manner with opposite effects at 9 days and 14 days. An RNAseq experiment reveals candidate genes potentially involved in this regulation. Our data highlight that sugar metabolism is intricately linked with other metabolite pathways. Alteration of single genes in central carbon metabolism profoundly alters plant immune responses and plant-microbe interactions.

## Introduction

Plants utilize sugars for growth and development: as carbon skeletons for enzymes and molecules, as energy source ensuring survival, as signaling compounds, and as currency to interact with beneficial and pathogenic microbes (Jeandet *et al*., 2022) Carbohydrates are produced in leaves via photosynthesis and transported to sink tissues. Belowground, plants are a source of reduced carbon for heterotrophic soil organisms. Carbon is supplied in the form of rhizodeposits, where root exudates comprise the soluble fraction. Exudates contain low- and high-molecular-weight compounds such as phenolics, amino acids, organic acids, terpenoids, lipids, nucleic acids, proteins, and sugars (Vives-Peris et al., 2019). Of the carbon fixed in leaves, between 5% and 30% is exuded (Finger and Möhring, 2024), mostly in the form of sugars and organic acids (Boeuf-Tremblay *et al*., 1995). Interestingly, exudation quantities and composition change with plant species, nutritional status, and development stage (Bais *et al*., 2006; Nakayama and Tateno, 2018). Differential exudation might be mediated by regulation of compounds biosynthesis, or by modulation of release from roots into the rhizosphere. In principle, metabolites can be exuded passively via (facilitated) diffusion or actively via energy-coupled transporter proteins. The few transporter proteins that have been characterized to be involved in exudation were shown to be critical regulators (Sasse *et al*., 2018). However, the mode of transport for most exuded compounds, including sugars, is unclear.

Various active sugar transporters have been described to be essential for sugar export from leaves, for loading and unloading of the phloem, and for sugar re-uptake from soil: among them are members of the SUC, STP, and SWEET families (Chen *et al*., 2015b; Doidy *et al*., 2012; Hennion *et al*., 2019). STP1 for example was the first plant hexose transporter reported to transport monosaccharides to sink tissues (Boorer *et al*., 1994; Sauer *et al*., 1990; Sherson *et al*., 2000). It is also involved in sugar re-uptake from the rhizosphere (Yamada *et al*., 2011). STP13 imports hexoses from the apoplasm into the cytosol in leaves. Interestingly, this transporter is activated by the immune kinase BIK1 upon pathogen perception to increase sugar uptake into cells, depleting apoplasm-located pathogens from energy (Yamada *et al*., 2016). Further evidence links sugar metabolism and immunity: exogenously applied sugars enhanced plant resistance to bacterial and fungal pathogens (Bolouri Moghaddam and Van Den Ende, 2012; Morkunas and Ratajczak, 2014). Moreover, elevated internal sugar levels increased salicylic acid synthesis, a key hormone involved in defense responses (Zhang and Li, 2019). However, how changes in sugar metabolism influence plant-pathogen and more generally plant-microbe interactions remains unclear.

Here, we characterize the role of sugar metabolism in plant-microbe interactions by studying *Arabidopsis thaliana* wild-type and sugar metabolism mutant lines, including starch-deficient mutants (*pgm1*, *adg1*, *pgi1*, *sex4* ), and an STP family sugar transporter (*stp1* ). We investigate i) general changes in metabolite profiles in mutant lines with altered sugar metabolism, ii) specific changes in hexose levels in tissues and exudates in sugar mutant lines, iii) altered immune and transcriptional responses of Col-0 when treated with external sugars, iv) altered responses of sugar mutant lines to pathogens, and v) changes in root-associated microbial numbers compared to Col-0. We describe an intertwined and complex interaction between sugar metabolism and immunity: Sugar levels impact the production of reactive oxygen species, an early immune response, as well as susceptibility to pathogens and interactions with root-associated microbial communities.

### Mutant lines with altered sugar homeostasis feature distinct tissue and exudate metabolite profiles

Sugars have a central role in plant metabolism. Thus, we hypothesized that plants with impaired starch and sugar metabolism not only feature altered monosaccharide and starch levels as reported (Streb and Zeeman, 2012) but also broad changes across many metabolite pathways. To test this, we selected a range of *Arabidopsis thaliana* (hereon: Arabidopsis) mutant lines devoid of starch biosynthesis (*pgm1*, *adg1*, *pgi1*), starch degradation (*sex4*), or hexose transport (*stp1*).

In a first step, we used a general metabolite profiling technique to assess changes in leaf, root, and exudate profiles due to altered carbon metabolism. In this untargeted metabolomic analysis of sterile-grown, three-week-old plants, 2,602 metabolites were detected in shoot and 2,617 metabolites in root tissues. The metabolite profiles collected at midday of several sugar metabolism mutant lines were distinct from Col-0 for shoots (*adg1*, *pgi1*, *stp1*, and *sex4*), and for roots (*adg1*, *pgi1*, and *stp1*, Figure 1 a and b).

**Figure 1:**
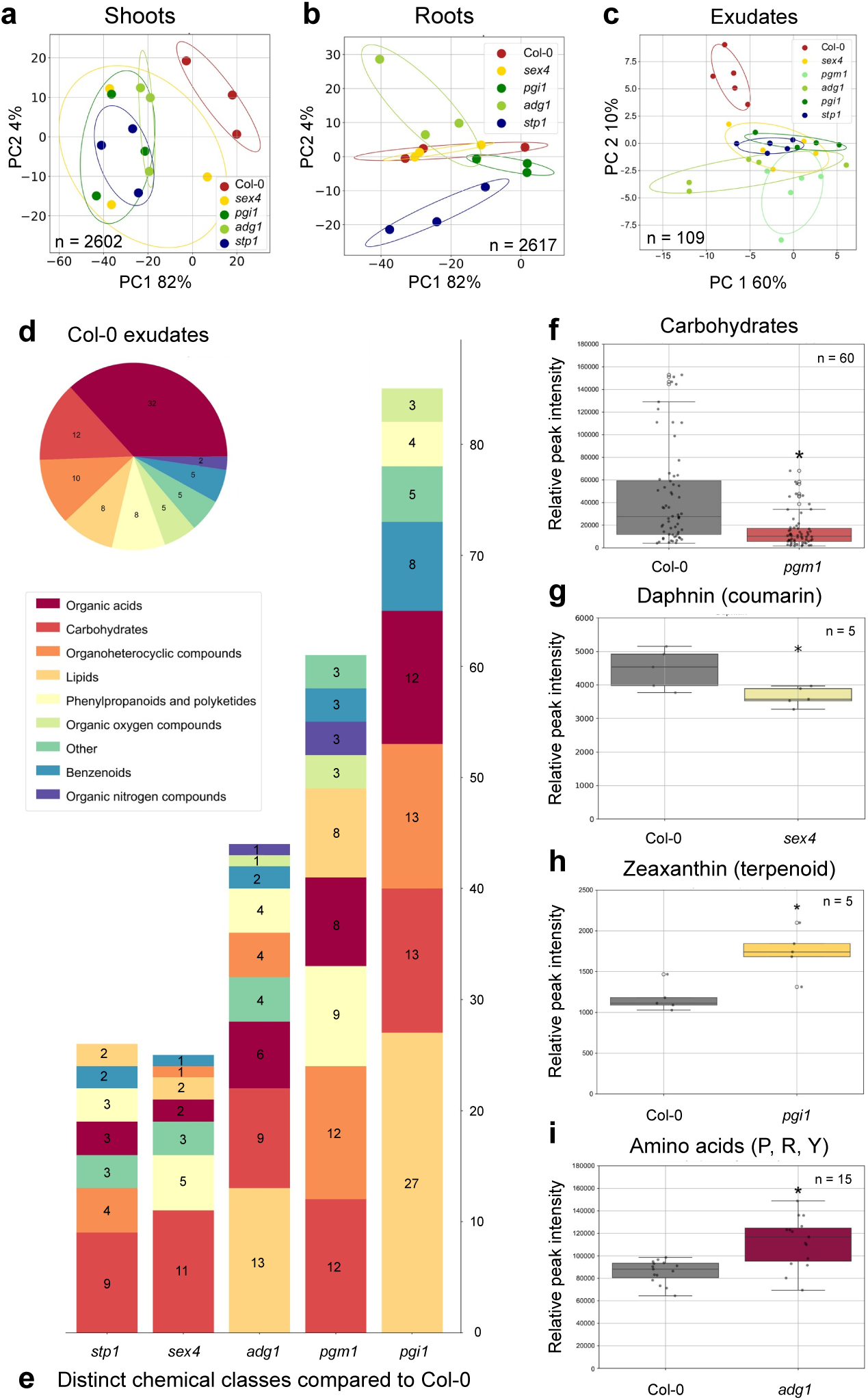
Sugar metabolism mutant lines feature broadly altered tissue and exudate metabolite profiles. (a-c) Principal component analysis (PCA) of metabolite profiles of shoots (a), roots (b), and exudates (c) of Col-0 and sugar metabolism mutant lines. The number of metabolites for the analyses is indicated in graphs (n). Variances of the PCA are expressed in percent in principal components (PC) 1 and 2. Data is based on 5 biological replicates (jars with each 5 Arabidopsis individuals). (d) Metabolites present in Col-0 exudates assigned to chemical superclasses (n=87) (Classyfire: https://cfb.fiehnlab.ucdavis.edu/, Djoumbou Feunang *et al*. (2016)). (e) Metabolites significantly distinct between exudates of Col-0 and sugar metabolism mutant lines categorized by chemical class. Colors: red: organic acids and derivatives, dark orange: carbohydrates and carbohydrate conjugates, orange: organoheterocyclic compounds, gold: lipids and lipid-like molecules, light yellow: phenylpropanoids and polyketides, green: Organic oxygen compounds, cyan: other (comprising non-assigned metabolites, lignans and neolignans, nucleosides and nucleotides, organophosphorus and organosulfur compounds, and alkanoids), petrol blue: benzenoids, purple: organic nitrogen compounds. Numbers represent the number of metabolites in each superclass. (f-i) Intensities of various metabolites compared to Col-0. (f) Carbohydrates in *pgm1*, (g) Daphnin (coumarin) in *sex4*, (h) Zeaxanthin (terpenoid) in *pgi1*, (i) Amino acids (P: proline, A: arginine, Y: tyrosine) in *adg1*. Number of replicates for each metabolite in the analyses is indicated in graphs (n). Data is displayed as average *±* S.E.M., *: pvalue < 0.05 (T-Test vs. Col-0).

In exudates collected for two hours midday, 109 metabolites were detected, which could be assigned to 8 chemical classes (Figure 1 c and d). In Col-0, organic acids and derivatives were most abundant, followed by carbohydrates and conjugates (also comprising sugars), organoheterocyclic compounds, lipids, phenylpropanoids, organic oxygen compounds, other organic biomolecules, benzenoids, and organic nitrogen compounds (Figure 1 d). Exudate profiles of all mutant lines were distinct from Col-0 on a principal component analysis (Figure 1 c): *pgi1* exudates were most distinct, featuring 85 compounds distinct from Col-0 (78%), followed by *pgm1* (65 metabolites/ 59.6%), *adg1* (44 metabolites/ 40.4%), *stp1* (26 metabolites/ 23.8%), and *sex4* (25 metabolites/ 23%) (Figure 1 e). For *pgi1*, most differences were found in lipids (27 metabolites, 31.8%), followed by carbohydrates (13 metabolites, 15.3%), organoheterocyclic compounds (13 metabolites, 15.3%), and others (Figure 1 e). Lipids were also differently exuded in exudates of *adg1*, and carbohydrates were majorly altered in exudates of all mutant lines (Figure 1 e).

Importantly, exuded carbohydrate levels were generally reduced in mutant lines compared to Col-0, as shown for *pgm1* (Figure 1 f). Other compounds such as coumarins, which are phenylpropanoids involved in defense responses, also decreased in *sex4* exudates (Figure 1 g), whereas other secondary metabolites and some amino acids increased in abundance (Figure 1 h and i). We conclude that sugar metabolism mutant lines exhibit broadly altered tissue and exudate metabolite profiles. As expected, the abundances of metabolites involved in carbohydrate metabolism were altered, but also the abundances of lipids, organic acids, and phenylpropanoids were affected.

### Sugar mutant lines exhibit reduced sugar exudation during the day

In a second step, we specifically quantified sugars in tissues and exudates to investigate how altered sugar metabolism affects their balance. For this, we used a specific chromatography technique as sugars are often not detected reliably with untargeted metabolomic approaches, as they do not ionize as efficiently as other metabolites, and as many sugars share their molecular mass.

Glucose, fructose, and sucrose levels were distinct in tissues of our sugar mutant lines compared to Col-0, as reported (Caspar *et al*., 1985; Lin *et al*., 1988a; Zeeman and Rees, 1999). In the middle of the day (MOD), root sugar profiles of the mutant lines were more distinct from Col-0 than shoot profiles (Figure 2 a and b, Suppl. Figure S.1). In addition, we report changes in raffinose and trehalose levels. In our dataset at midday, glucose levels were not significantly altered in any mutants or tissues. In shoots, raffinose levels were decreased in *pgm1* and *pgi1* compared to Col-0, and sucrose was elevated in *adg1* (Figure 2 a). The 70% raffinose decrease in *pgm1* shoots was the largest change observed in the dataset. In roots, sucrose, fructose, and raffinose accumulated in *pgm1* (Figure 2 b). Overall, *pgm1* was the mutant line strongest affected in sugar levels in both root and shoots, while the other mutants only exhibited changes in one type of tissue.

**Figure 2:**
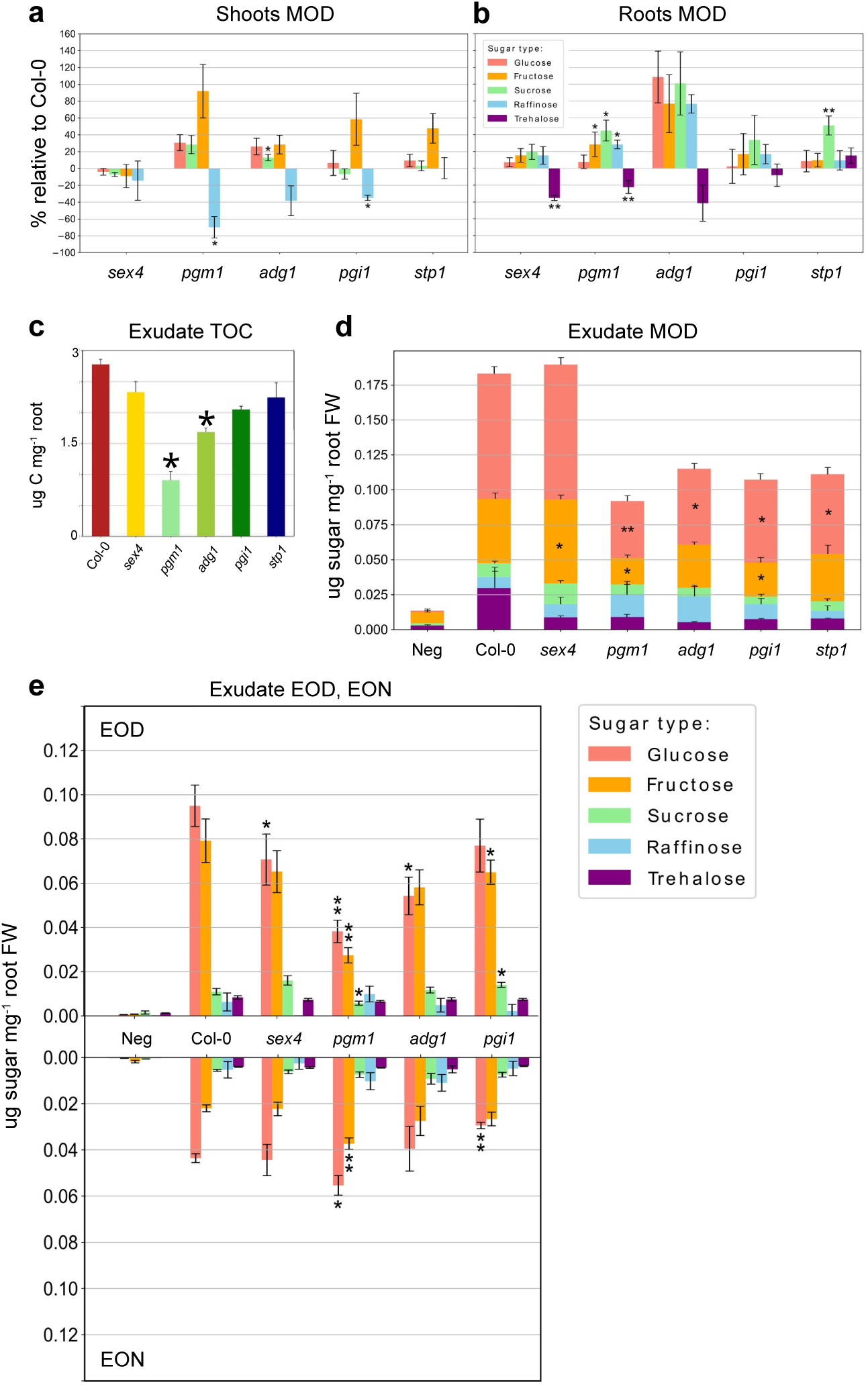
Sugar metabolism mutant lines exhibit altered sugar exudation. (a-b) Middle of day (MOD) shoot (a) and root (b) sugar content of *sex4*, *pgm1*, *adg1*, *pgi1*, and *stp1* presented in percent relative to Col-0 (positive values: higher than Col-0, negative values: lower than Col-0). Data is displayed as average *±* S.E.M., *: pvalue < 0.05, **: pvalue < 0.01 (T-Test vs. Col-0 corresponding sugar). For non-relative sugar content of tissues, see Suppl. Figure S.1 (c) Total organic carbon (TOC) determination of Col-0 and sugar metabolism mutant exudates. Data is displayed as average *±* S.E.M., *: pvalue < 0.05 (T-Test vs. Col-0). (d) MOD exudate sugar content of Col-0 and sugar metabolism mutant lines as well as exudate negative controls (no plant present). Data is based on 5 biological replicates (jars with each 5 Arabidopsis individuals), and displayed as average *±* S.E.M., *: pvalue < 0.05, **: pvalue < 0.01 (T-Test vs. Col-0 corresponding sugar). (e) End of day (EOD, positive y-axis) and end of night (EON, negative y-axis) exudate sugar content of Col-0 and sugar metabolism mutant lines as well as exudate negative controls (no plant present). Data is based on 5 biological replicates (jars with each 5 Arabidopsis individuals), and displayed as average *±* S.E.M., *: pvalue < 0.05, **: pvalue < 0.01 (T-Test vs. Col-0 corresponding sugar). Colors for different sugars: pink: glucose, orange: fructose, green: sucrose, blue: raffinose, purple: trehalose. For raw data on sugar content in tissues and exudates see Suppl. Figure S.2.

We hypothesized that our sugar metabolism mutant lines would feature altered sugar exudation: Starch mutant lines exhibit increased soluble sugar levels during the day and starve during the night, resulting in smaller plant size overall (Martin and Smith, 1995). Thus, we hypothesized that sugar exudation may be increased during the day and reduced during the night. We first assessed exudation quantity by determining total organic carbon (TOC). In contrast to our hypothesis, *pgm1* and *adg1* featured lower carbon exudation levels compared to Col-0 at midday (Figure 2 c).

Utilizing the sugar-specific chromatography method, all sugars quantified in tissues could also be detected in exudates (glucose, fructose, sucrose, trehalose, raffinose, Figure 2 d, Suppl. Figure S.2). The most abundant sugar exuded in Col-0 midday was glucose (90 ng mg^−1^ root) followed by fructose (45 ng mg^−1^ root), trehalose (30 ng mg^−1^ root), sucrose (10 ng mg^−1^ root), and raffinose (8 ng mg^−1^ root) (Figure 2 d). Compared to Col-0, *pgm1*, *adg1*, *pgi1*, and *stp1* exuded 33-55% less glucose, with the strongest reduction observed in *pgm1*. Fructose exudation was similarly reduced in *pgm1* and *pgi1* by 61% and 48%, respectively. Interestingly, *sex4* exuded 28% more fructose compared to Col-0.

It is interesting to note that compared to tissues (Figure 2 a and b), in which glucose levels remained unchanged between lines, the exuded glucose content was decreased for all biosynthesis mutants and the sugar transporter mutant (Figure 2 d). Further, although sugars accumulated in *pgm1* roots (Figure 2 b), glucose and fructose levels were decreased in exudates (Figure 2 d). Similarly, although *sex4* sugar levels in tissues were comparable to Col-0 (Figure 2 a and b), fructose exudation was increased (Figure 2 d). The reverse observation could also be made: although *pgm1* and *stp1* showed alterations in root sugar content (Figure 2 b), these changes were not reflected in exudation (Figure 2 d). Thus, we conclude that sugar changes observed in tissues are not directly reflected in exudates, pointing towards a non-passive mode of sugar exudation.

In summary, we show with three different techniques, untargeted metabolomic profiling, TOC, and a sugar-specific chromatography, that sugar levels are decreased in exudates during the day when sugar metabolism in plants is altered. In most lines, night-time exudation remained unchanged, but in *pgm1*, an increase was observed.

### Sugar exudation is higher during day than night, except for *pgm1*

To further explore exudate dynamics in starch mutant lines, exudates were collected for two hours at the end of the day (EOD) and the end of the night (EON), and sugar levels were quantified (Figure 2 e, Suppl. Figure S.2). For Col-0, the total exudation was at 0.2 µg mg^−1^ root (0.008 µg mg^−1^ trehalose, 0.096 µg mg^−1^ glucose, 0.011 µg mg^−1^ sucrose, 0.079 µg mg^−1^ fructose, 0.006 µg mg^−1^ raffinose) at the EOD and 2.5-fold lower at the EON (0.08 µg mg^−1^ root). Generally, EOD sugar levels were comparable to MOD levels. Glucose, as observed previously, was the most abundant in all samples, and was the most affected in the mutant lines. Glucose and fructose levels were reduced in *pgm1*, and glucose was reduced in *adg1* compared to Col-0 at the EOD (−59,8% and −42.9% respectively, Figure 2 e). In contrast to the MOD, *sex4* showed a significant reduction in glucose exudation EOD and *pgi1* only showed a significant drop in glucose exudation at the MOD and the EON but not at the EOD. These observations suggest that sugar exudation dynamics are complex, and more thorough investigations are warranted to resolve the diurnal dynamics of tissue and exudate sugar levels. Generally, exudation levels of *sex4*, *adg1*, and *pgi1* followed the same trend with higher sugar levels at EOD compared to EON. Paradoxically, *pgm1* featured an opposite pattern, releasing approximately 30% more sugars at the EON compared to the EOD, implying that different starch biosynthesis mutations do not necessarily have the same effect on root exudation.

### External sucrose levels alter ROS response depending on developmental stage

In a first effort to investigate how sugar metabolism influences plant immune responses, we monitored the production of reactive oxygen species (ROS) and early immune response. In typical ROS assays with young plants, they are grown with 1% sucrose in liquid culture for 9 d (Bjornson *et al*., 2021). To investigate whether the external sugar supply impacts ROS production, Col-0 plants were grown on solid MS plates supplied with or without 1% sucrose for 9 d, 14 d, or 21 d. The plants were then treated with flg22, elf18, or pep1, typical elicitors resulting in ROS production (Felix *et al*., 1999; Kadota *et al*., 2015; Kunze *et al*., 2004).

For 9 d-old seedlings, the immune response was higher when supplied with external sucrose (Figure 3 a and d). Interestingly, this pattern was reversed in 14 d-old plants, in which the immune response was lower with external sucrose present (Figure 3 b and d). At 21 d, plants with and without external sucrose supply exhibited the same response (Figure 3 c and d). This assay was repeated with sugar mutant lines, which showed comparable patterns to Col-0 (Suppl. Figure S.4 a-o). This data implies that the addition of sucrose influences the ROS immune response and that this response is dependent on the plant developmental stage but is not shaped by the mutations in sugar and starch pathways investigated.

**Figure 3:**
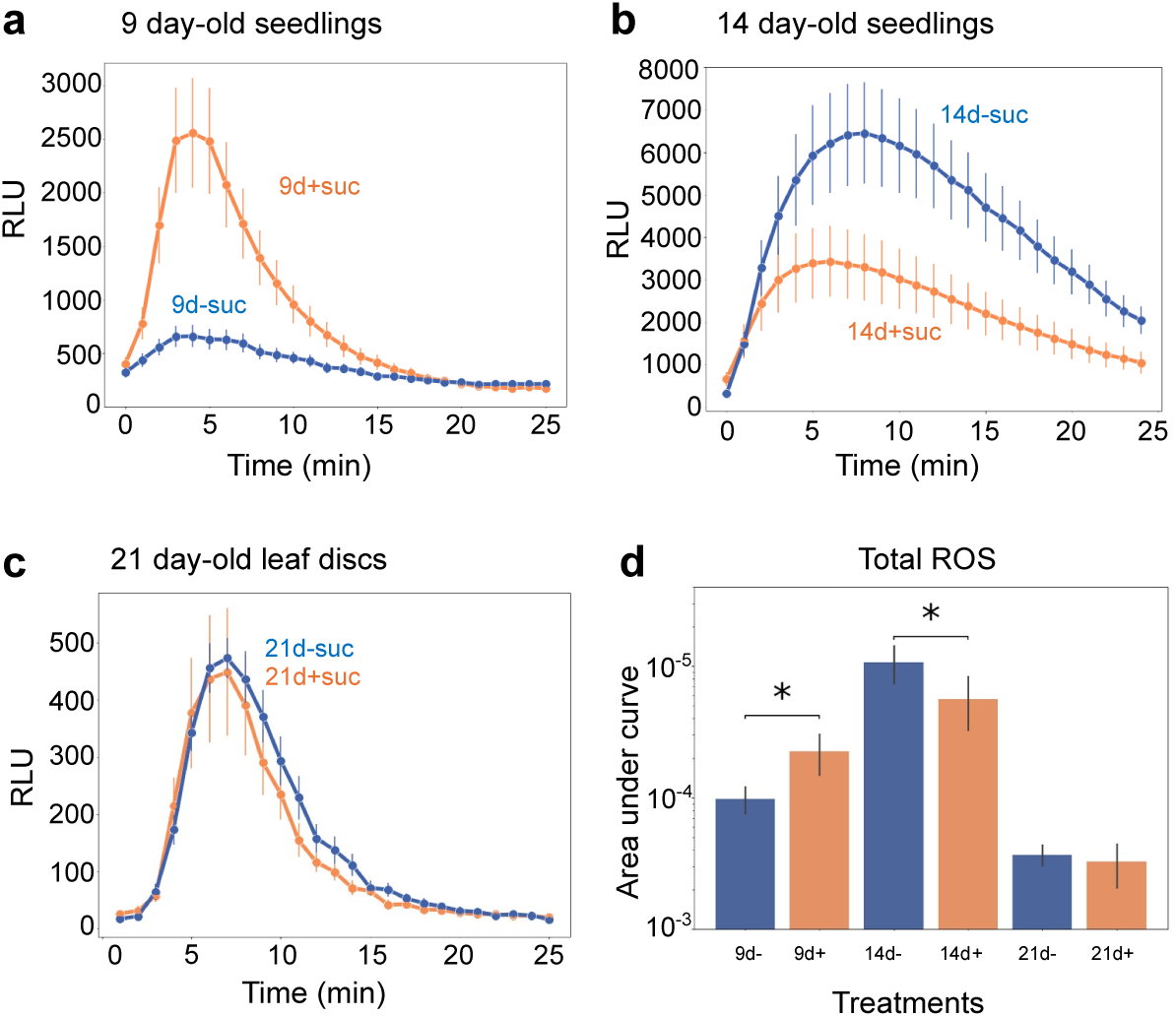
Reactive oxygen species production depends on presence of external sucrose and plant developmental stage. (a-c) Reactive oxygen species (ROS) production for 9 d (a), 14 d (b), and 21 d-old Col-0 (c). Plants were grown on ½ MS (Murashige and Skoog) plates with or without supplementation of 1% sucrose (orange and blue curves, respectively) and treated with 100 nM flg22. (d) Area under the curve for all samples in a-c. Data is based on 8 replicates for 9 and 14 d-old seedlings and 4 replicates for 21 d-old leaf discs. Data is displayed as average *±* S.E.M., *: pvalue < 0.05 (T-Test with vs without sucrose samples). For ROS burst of Col-0 upon elf18 and pep1 elicitor treatments see Suppl. Figure S.3. For ROS burst upon flg22 elicitor treatment (100 nM) of sugar metabolism mutant lines see Suppl. Figure S.4. +: with sucrose, −: without sucrose. RLU: relative light units.

### Sucrose supplementation results in elevated expression of defense-related genes at 9 d and of development-related genes at 14 d

We wanted to determine the genetic basis for the opposite ROS production when supplied with external sucrose (Figure 3 a-d). We conducted an RNAseq experiment for Col-0 grown with and without 1% external sucrose. Plants were harvested at 9 d, 14 d, and 21 d, and gene expression patterns were investigated (Figure 4 a). We hypothesized that filtering differentially expressed genes correlating with the observed ROS phenotype would result in list of candidate genes causing the distinct response. Overall, 33,610 genes were expressed in the dataset. Of these, 9.7% (3,261 genes) of genes were significantly distinct between sucrose treatments at 9 d, 11.2% (3,761 genes) at 14 d, and 3.3% (1,101 genes) at 21 d. Clustering of differentially expressed genes of all time points revealed four clusters (Figure 4 a). Consistent with the ROS phenotype, young plants exhibited a strong differential gene expression, whereas old plants did not (9 d and 14 d vs 21 d). The largest cluster, cluster II, comprised genes that were upregulated in response to sucrose in both 9 d and 14 d-old plants. A subset of cluster II showed upregulation of gene expression in response to sucrose for all differential aged plants. The second-largest cluster IV exhibited opposite regulation to cluster II with genes that were downregulated in response to sucrose in both earlier time points. Cluster I and III comprised genes that showed an opposite regulatory pattern in response to sucrose at 9 d and 14 d. In cluster I, the addition of sucrose triggered the downregulation of genes in 9 d and upregulation in 14 d old plants, whereas plants at 21 d did not exhibit differential expression in response to sucrose. Cluster III showed the opposite pattern compared to cluster I. Thus, cluster I and III potentially present interesting candidates to explain the ROS phenotype observed.

**Figure 4:**
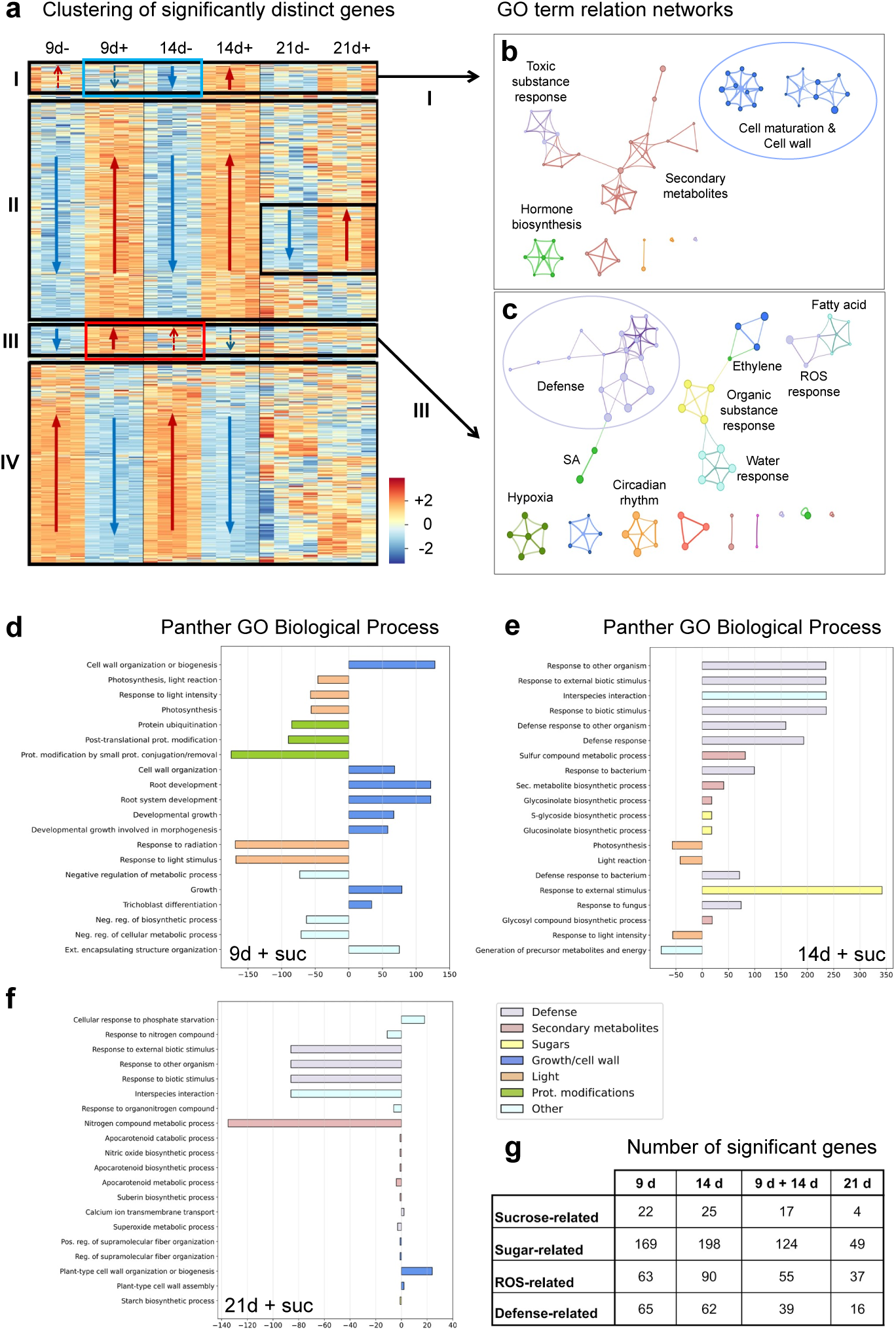
External sucrose elicits early defense-related and late development-related genes. (a) Hierarchical clustering of differentially expressed genes in a heatmap from 9 d, 14 d, and 21 d-old Col-0 grown with (+) or without (−) sucrose addition. Color code: red: positive 2-fold change and upregulation, blue: negative 2-fold change and downregulation. Data is based on 4 biological replicates. (b-c) Metascape gene ontology (GO) relation network of cluster I (b) and cluster III (c) genes (Zhou *et al*., 2019). Colors: blue: cell growth process, light purple: defense-related process, pink: secondary metabolites, green: hormones, orange: time-dependent process, yellow: organic substance response, olive green: hypoxia response, red: signal transduction process, magenta: iron ion transport, light blue: others (water, fatty acids processes). (d-f) 20 most significant PANTHER GO biological processes based on the comparison of samples grown with and without sucrose for 9 d (d), 14 d (e), and 21 d (f). Colors: light purple: defense, pink: secondary metabolites, yellow: sugars, blue: growth and cell wall development, orange: light, green: protein modifications, light blue: others. (g) The number of genes in 4 manually created lists of genes related to sucrose, sugars, ROS, and defense built from the significant numbers of metabolites in 9 d, 14 d, and 21 d-old plants.

Clusters I and III were first analyzed with a Metascape gene ontology (GO) network analysis (Figure 4 b and c, Zhou *et al*. (2019)). the cluster I network comprised functions related to growth, such as cell maturation, cell wall expansion, and hormone biosynthesis. In contrast, the cluster III network comprised functions related to defense, such as ROS response, hypoxia, salicylic acid, and ethylene responses. A second analysis with the PANTHER tools was based on the full set of differentially expressed genes upon sucrose addition (Figure 4 d-f, Mi *et al*. (2019)). Several interesting patterns were observed: i) At 9 d, sucrose addition resulted in upregulation of 16/20 of the most significant GO terms. Eight upregulated terms were related to responses to biotic stimuli/defense, and four more to secondary metabolite pathways. Downregulated pathways were related to photosynthesis and energy metabolism (Figure 4 d). ii) At 14 d, only half of the most significant GO terms showed upregulation, and most of these terms were related to development and growth. Moreover, light and protein modifications were strongly decreased (Figure 4 e). iii) at 21 d, sucrose addition resulted in less pronounced gene regulation overall (Figure 4 f).

Next, to determine gene candidates potentially explaining the observed phenotype, we methodically attributed significant genes to the four following categories by browsing gene descriptions and literature: sucrose-related genes (GO term sucrose response), sugar-related genes (genes of carbohydrate and starch pathways), ROS-related genes (Mittler *et al*., 2004), and defense-related genes (immune-related genes, such as receptor proteins, Figure 4 g). Most of the genes of all four lists were similarly regulated at 9 d and 14 d (Figure 4 g). For example, for sucrose-related genes, 22 were differentially regulated at 9 d and 25 genes at 14 d, with similar regulation of 17 genes across both time points (Figure 4 g). Thus, the regulation of these manually selected genes reflected the data of the heatmap, where most of the genes were similarly regulated also (Figure 4 a, clusters III and IV). Interestingly, in the ROS-related genes list, RBOHD (AT5G47910) showed up as being upregulated for sucrose supplemented 9 d and 14 d but not for 21 d. Similarly, in the defense-related genes list, defense receptors (FLS2, PR1-like, RLP30) were upregulated in sucrose supplemented 9 d and 14 d, but not in 21 d). Moreover, we focused on the few genes with opposite expression patterns in clusters I and III (Figure 4 a) to investigate the genetic cause for the differential ROS response. We found members of several transcription factor families differentially regulated in clusters I and III that might coordinate the response (bHLH, ERF, MYB, WRKY). Members of b-glucosidases (BGLU5 11, 21, 22, PKY10), of the recently characterized Ca^2+^ Channels (MLO12, MLO15), and of immune- and defense-related genes (RLP23, UGT73C7, CAF1b, LOX4, NUDT7, MKK4) might link carbohydrate- and immune-regulated reponses. Interestingly, several amino-acid-related genes and transporters (AAP6, UmamiT4, 12, 21, 37, 46, GDU5, LHT1) were detected in clusters I and III, potentially highlighting a role for these metabolites in mediating response to extracellular sugar. Last, auxin-related genes (YUC3, 4, 5, 9, TAA1, AXR3, SAUR9, 26, 28, NIT2) were differentially regulated, which might mediate the development-related responses to extracellular sugar. In summary, we found the expression of defense-related genes to be increased, and of development-related genes to be decreased, which might explain the differential ROS response to external sucrose.

### Reduced pathogen levels in sugar mutant lines

As we found many defense-related genes to be differentially regulated in the RNAseq experiment, and as sugars are nutrients to pathogens and signals to plant immunity, we next investigated whether sugar metabolism mutant lines were responding differently to pathogen presence. For this, Col-0 and mutant lines were infected with *Pseudomonas syringae* DC3000, a leaf pathogen. Compared to Col-0, infection levels were higher in the elicitor-insensitive control as expected, and lower in *sex4*, *pgi1*, and *stp1* (Figure 5 a). This data suggests that some sugar metabolism mutant lines are more resistant to leaf pathogens, which could be caused by an increase in immune responses, or by relocation of sugars to compartments inaccessible for the pathogen. Interestingly, the increased pathogen response was not directly correlated with leaf sugar content measurements (Figure 2 a): *sex4* and *stp1* for example featured increased pathogen resistance but had overall comparable sugar levels compared to Col-0. The *pgi1* exhibited lower raffinose levels as did *pgm1*, but whereas *pgi1* featured increased pathogen resistance, *pgm1* did not. To resolve this, sub-cellular sugar allocation should be investigated in these sugar metabolism mutant lines. We conclude that although some sugar mutant lines exhibit elevated pathogen resistance, this is not directly correlated with glucose, fructose, sucrose, or raffinose levels in leaves.

**Figure 5:**
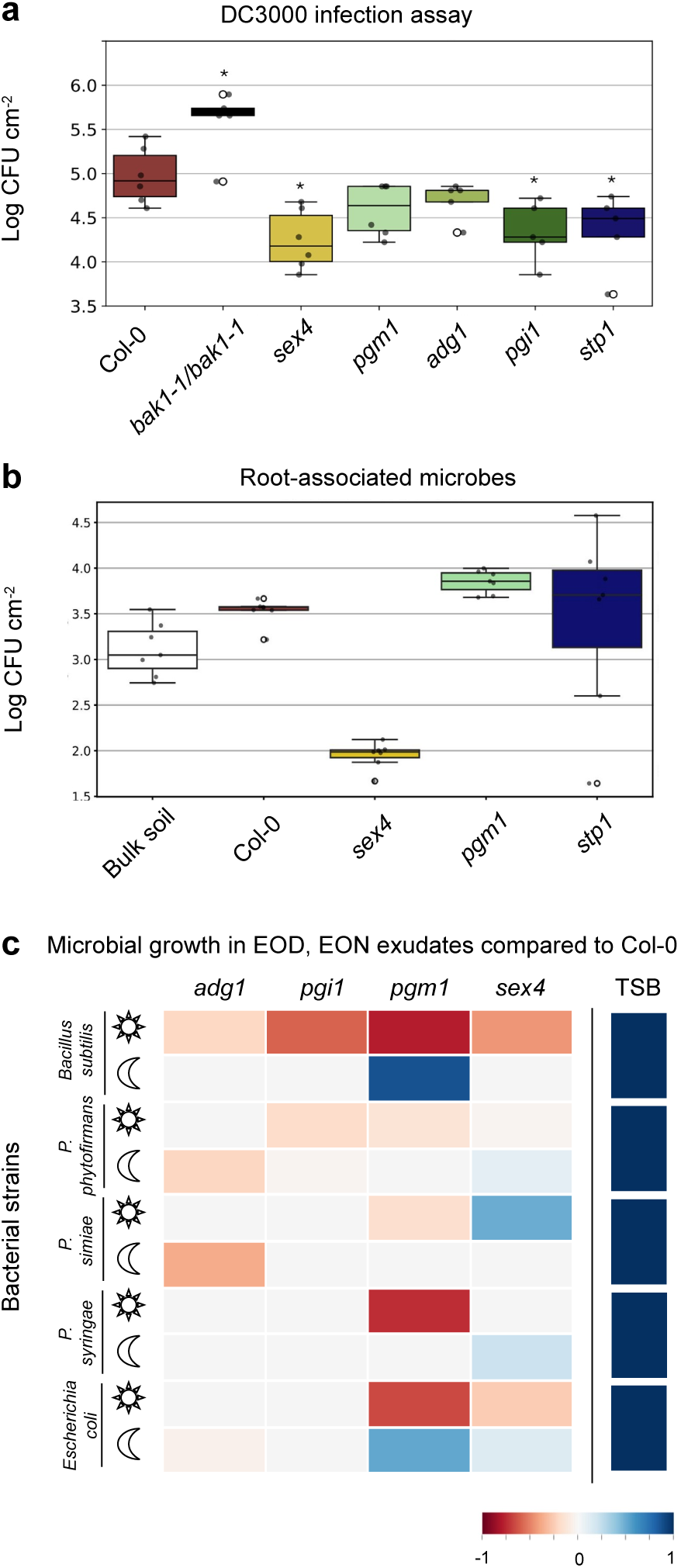
Altered sugar metabolism affects plant-microbe interactions. (a) Pathogen infection assay of 3-week-old Col-0 (red), bak1-1/bak1-1 (black, negative control, insensitive to pathogen), *sex4* (yellow), *pgm1* (cyan), *adg1* (light green), *pgi1* (forest green), *stp1* (navy blue) with *Pseudomonas syringae* pv tomato DC3000. Data is displayed as colony forming units (CFU) per leaf area. Data is displayed as average *±* S.E.M., *: pvalue < 0.05 (T-Test vs. Col-0), n =6 biological replicates. (b) Counts of root-associated microbes on Col-0 (red), *sex4* (yellow), *pgm1* (cyan), *stp1* (navy blue) for 21 d-old plants grown in standard soil. Bulk soil: soil samples collected from pots without plants. Data is displayed as average *±* S.E.M., *: pvalue < 0.05 (T-Test vs. Col-0), n = 8 biological replicates. (c) Bacterial growth on concentrated exudates collected EOD and EON of *adg1*, *pgi1*, *pgm1*, and *sex4* displayed as a heatmap of the area under the curve relative to growth in Col-0 exudates. Tryptic soy broth (TSB) was used to confirm bacterial activity. Microbes: *Bacillus subtilis*, *Paraburkholderia phytofirmans*, *Pseudomonas simiae*, *Pseudomonas syringae* DC3000, *Escherichia coli*. Color code: grey: no change in growth in comparison to Col-0, dark blue to light blue: higher growth in comparison to growth on Col-0 exudates, dark red to light red: decreased growth in comparison to growth in Col-0 exudates. For the heatmap of the area under the curve relative to growth in Neg (empty jar) exudates, see Suppl. Figure S.5.

### Altered microbial growth on sugar mutant exudates

As changes in sugar metabolism affected plant-pathogen interactions, we next investigated whether other plant-microbe interactions were affected as well. For this, we investigated how many soil-derived microbes associate with roots, and how single microbial strains performed when grown in exudates of various sugar metabolism mutant lines.

Col-0 and sugar metabolism mutant lines were grown in soil for three weeks, and root-associated microbes were quantified by counting colonies originating from the rhizosphere (Figure 5 b). Col-0 root-associated microbial counts were slightly higher than in the bulk soil. Interestingly, *pgm1* roots exhibited higher, and *sex4* lower colonization levels compared to Col-0. As in *pgm1*, sugar daytime exudation was lower and nighttime exudation higher compared to Col-0 (Figure 2 e), this increase in root-associated microbes could be due to the elevated EON sugar levels or alternatively, by the distinct exudate composition observed (Figure 1 e). The lower colonization of *sex4* is likely not caused by altered sugar exudation, as levels were comparable to Col-0 (Figure 2 e) but might be caused by changes in e.g. other carbohydrates, or phenylpropanoids, (Figure 1 e).

To further test the hypothesis of whether altered exudation of sugars and other compounds resulted in differential microbial growth, phylogenetically distinct microbes were grown in concentrated exudates collected at the EOD and the EON, and their optical density (OD600) was monitored over time (Figure 5 c). All bacterial strains showed growth on exudate samples compared to negative samples and in a complex nutrient medium (Suppl. Figure S.5). Additionally, we confirm that they could all grow well in our setup when supplied with a high nutritive medium (TSB, Figure 5 c). *Bacillus subtilis*, *Paraburkholderia phytofirmans*, and *Pseudomonas simiae* were selected for their potentially beneficial effect on plant health (Rajkumar *et al*., 2017; Tiwari *et al*., 2019). All of these microbes grew less efficiently on exudates of sugar metabolism mutant lines compared to Col-0 exudates if there was a significant difference, except for *P. phytofirmans* and *P. simiae* growing on *sex4* exudates and *B. subtilis* growing on *pgm1* EON exudates. Further, we also tested the pathogen *P. syringae* DC3000 that was used for the leaf infection assay and *Escherichia coli*, which is not specifically associated with plants. Both microbes grew less efficient in EOD exudates of *pgm1* and better on *sex4* EON exudates, but else not different from Col-0 exudates. Interestingly, this is the opposite pattern as observed for the root-associated microbiome, which was higher in *pgm1* and lower in *sex4*. Taken together, the strongest microbial growth effects were seen in *pgm1* exudates, with EOD exudates consistently featuring lower bacterial growth, likely caused by limited sugar exudation during the day (Figure 2 e). Consistently, *pgm1* EON exudates increased the growth of 2 out of 5 bacteria, indicating that the differential sugar exudation observed might indeed impact root-microbe interactions. We conclude that altered exudation profiles in general and altered carbohydrate/TOC exudation indeed impacts microbial growth. In a next step, it would be important to determine the exuded metabolites altering growth of specific strains.

## Discussion

Sugars are key compounds in plant metabolism, growth, and development, and interact closely with the plants immune system as signals. Here, we show that alterations in sugar metabolism, either by the addition of external sugar or by impairing genes of starch and sugar metabolite pathways, result in complex changes not only in sugar levels, but also in many other metabolite pathways across shoots, roots, and exudates (Figures 1 and 2).

Effects of altered starch metabolism and transport have been thoroughly investigated in leaves (Büttner, 2007; Streb and Zeeman, 2012). Glucose, fructose, and sucrose levels fluctuate throughout the day and night (Chia *et al*., 2004). Although our plants were grown in a sterile, semi-hydroponic system and a different method was used for sugar quantification, our leaf sugar values align with values published: at midday, our leaves contained 1.24-1.5 µg mg^−1^ leaf fresh tissue of glucose, 0.4-0.45 µg mg^−1^ fructose, and 0.31-0.7 µg mg^−1^ sucrose (Suppl. Figures S.1 and S.2, Chia *et al*. (2004)). In contrast to leaves, sugar metabolism in roots and exudates is far less characterized. Few studies quantify sugar levels. In tomato roots, total soluble sugar levels (glucose, fructose, sucrose) fluctuated between 20-30 µg mg^−1^ root dry weight (Devaux *et al*., 2003). In Arabidopsis Col-0, the levels of glucose and fructose are similar in both roots and shoots. However, the levels of sucrose, raffinose, and trehalose are up to 3 times higher in the roots compared to the shoots. In contrast, sugar levels are not comparable between leaves and roots for all sugars and all mutants. For example, Glucose contents in shoots and roots are similar except for *adg1*. Fructose is also mostly similar between shoots and roots except for *pgm1* and *pgi1*. In roots, sucrose is 2 to 5 times higher than in shoots. Raffinose is 3 to 7 times higher in roots than in shoots. Finally, trehalose is only present in roots. The altered root sugar levels might impact root morphology and microbial associations. Indeed, several studies suggest that carbon availability in roots shapes root morphology and local hexose concentrations likely have a signaling function (Freixes *et al*., 2002; Singh *et al*., 2014). We further quantified sugars in exudates: Col-0 exudates contained 10-20 fold lower glucose, fructose, and sucrose levels compared to leaves.

We further quantified raffinose and trehalose in tissues and exudates. Both sugars are typically 10 to 100-fold less abundant in exudates compared to glucose and fructose, the soluble sugars with the highest abundance. Raffinose is typically induced under abiotic stress such as cold, drought stress, or oxidative stress (Li *et al*., 2020; Nishizawa *et al*., 2008), and trehalose regulates ROS accumulation, turgor pressure, and mineral balance by activating molecular and metabolic components involved in these processes, such as improvements of photosynthetic activity, stimulation of antioxidants, and effects on the expression of specific stress-response genes among others (Raza *et al*., 2023). Raffinose and trehalose were only changed in single mutant lines and tissues, but the method presented is a promising avenue for future research to determine the role of these sugars in plant development and plant-microbe interactions.

In addition to the ubiquitous quantification of sugars in tissues, we report sugar quantification in root exudates. Exudates are nutrients and signals to the microbes and animals present in soil (Sasse *et al*., 2018). Thus, analysis of qualitative and quantitative changes in exudates is essential for understanding pathogenic and beneficial plant-microbe interactions. Exudation profiles of sugar metabolism mutant lines are generally shifted, and feature altered sugar exudation. *pgm1* exudates were most distinct from Col-0 wild type, with a 50% reduction of total organic carbon and sugar exudation during the day, coupled with increased sugar exudation during the night. Further, *pgm1* featured altered carbohydrates, phenylpropanoid, organoheterocyclic compound, and organic acid exudation. Daytime-collected *pgm1* exudates consistently supported less growth of a variety of microbes in monoculture, whereas nighttime-collected exudates supported growth of *Bacillus subtilis* and *Escherichia coli*. *pgm1* lines feature comparable net photosynthesis rates to Col-0 wild type, elevated shoot-root assimilate transport, and increased root respiration over the course of 24 h (Brauner *et al*., 2014). Our data indicate that the diurnal exudation pattern for *pgm1* is altered. To characterize the molecular mechanisms resulting in altered interactions with root-associated microbes, a time-resolved carbon balance study should be made, including measurement of photosynthesis, respiration, carbon partitioning, and exudation.

We showed that external sugar application alters reactive oxygen production in a development-dependent manner. Whereas in 9-d-old plants ROS production was increased upon sucrose supplementation, 14-day-old plants exhibited a decrease, and 21-d-old plants exhibited the same ROS response independent of sucrose addition. An RNAseq analysis highlighted sugar-dependent transcriptional upregulation of defense-related genes and downregulation of development-related genes at 9 d upon sucrose addition, likely explaining the high ROS production. Generally, the candidate genes identified here as correlating with the ROS phenotype observed should be confirmed in future studies.

We showed that multiple sugar metabolism mutant lines have altered interactions with pathogens and soil-derived microbes. This data further supports the hypothesis the exudation shapes the microbiome (Huang *et al*., 2019; McLaughlin *et al*., 2023b; Pacheco-Moreno *et al*., 2024; Rudrappa *et al*., 2008; Sasse *et al*., 2018). Sucrose, trehalose, and jasmonic acid (JA) exudation were shown to shape the rhizosphere bacterial community composition in maize (Lopes *et al*., 2022). Compellingly, sucrose exudation had prominent effects early in development and trehalose at later stages. We observe distinct effects of external sucrose on plant immunity, and sugar exudation is predominant in early developmental stages across plant species (Clayton *et al*., 2008). Thus, it would be crucial to repeat the presented experiments at different developmental stages to resolve in a detailed manner how sugars shape plant-microbe interactions.

The importance of sugar transporters in plant-microbe interactions was investigated here by examining *stp1*, which exhibited altered exudation of organoheterocyclic compounds, carbohydrates and others, and showed increased resistance against *P. syringae*. Leaf-expressed STP13 is activated upon pathogen perception by the immune system kinase BIK1 to remove hexoses from the apoplasm (Yamada *et al*., 2016). If a similar mechanism exists for STP1 remains to be determined. SWEET sugar transporters are central to pathogen resistance in various tissues and species (Breia *et al*., 2021, 2020; Chen *et al*., 2015a, 2012; Yao *et al*., 2022). Recently, SWEET transporters were also implicated in shaping root-associated microbiomes in different root zones (Loo *et al*., 2024). We conclude that not only sugar metabolite genes but also sugar transporters are essential for plant-microbe interactions.

To summarize, we show that sugar metabolism, exudation, plant immune responses, and interactions with pathogenic and soil-derived microbes are intricately linked. We present methods suited for analysis of low concentrations of sugars in tissues and exudates. This work can be expanded towards a description of a full carbon balance in plants, in which carbon fluxes can be accurately modeled and engineered. We highlight that altered leaf sugar metabolism impacts carbon partitioning, exudation, and above- and belowground plant-microbe interactions. We advocate for a more holistic approach when studying aspects of plant metabolism and plant-organism interactions to account for this interconnectedness.

## Acknowledgments

We thank Prof. Dr. Nicola Zamboni (ETH Zurich, Switzerland) for acquisition of metabolite profiles of tissues and exudates, Prof. Dr. Samuel Zeeman and Dr. Michaela Fisher-Stettler (ETH Zurich, Switzerland) for sugar quantification and experimental support on HPAEC-PAD, Prof. Dr. Prof Klaus Schläppi and Charlotte Joller (University of Basel, Switzerland) for providing support and instrumentation for the bacterial growth assay. We thank Lay Sprecher for experimental support and Dr. Eva Marina Stirnemann for performing the RNA extraction, Prof. Dr. Leo Eberl (University of Zurich, Switzerland), Prof. Dr. Zeeman, and Prof. Dr. Cyril Zipfel (University of Zurich, Switzerland) for provision of seeds and microbial strains. We thank Prof. Dr. Zipfel and Prof. Dr. Niko Geldner for discussions. Finally, we acknowledge the Swiss National Science Foundation (PR00P3_185831 to J.S., supporting A.S. and J.S.).

## Materials and methods

### Plant material

Experiments were performed with *Arabidopsis thaliana* Columbia-0 (Col-0), *pgm1* (At5g51820, CS210), (Caspar *et al*., 1985), *sex4-3* (At3g52180, Salk_126784C), (Kötting *et al*., 2009; Zeeman and Rees, 1999), *adg1-1* (At5g48300, Salk_059083C), (Lin *et al*., 1988a,b; Wang *et al*., 1998), *pgi1* (At4g24620, Salk_107903C), (Bahaji *et al*., 2015), *stp1* (At1g11260, Salk_048848C), (Cordoba *et al*., 2015; Yamada *et al*., 2016), and *stp4* (At3g19930, Salk_091229C), (Poschet *et al*., 2010; Yamada *et al*., 2011).

### Sterilization, germination, and growth conditions

*Arabidopsis thaliana*seeds were sterilized for 15 min in 70% ethanol, and for 15 min in 100% ethanol, and air-dried in sterile conditions for 3 h. Seeds were plated on half-strength Murashige and Skoog medium (1/2 MS, M0221.0050, Duchefa Biochemie, Haarlem, Netherlands) supplied with 0.7% phytoagar (P10003.1000, Duchefa Biochemie, Haarlem, Netherlands), adjusted to pH 5.7 with KOH. Seeds were stratified for two days at 4°C in the dark, and grown in a 16 h light (150-160 mmol m^−2^ s^−1^) 22°C /8 h dark 18°C environment.

### Semi-hydroponic setup for metabolite analysis

The semi-hydroponic system was prepared in sterile conditions as published (McLaughlin *et al*., 2023a). Briefly, glass jars with glass beads were sterilized and supplied with liquid 1/2 MS medium. Sterility of the system was confirmed by plating a 20 µl aliquot on solid LB medium followed by incubation for 48 h at 23°C and monitoring for microbial growth. Five biological replicates (jars) each consisting of 5 plants were prepared. Negative controls comprised of 5 replicates (jars) to capture the metabolite background of the respective systems.

### Metabolite sample collection and analysis

The growth medium was exchanged with 50 ml of filter-sterilized 20 mM ammonium acetate, pH 5.7 (102308585, Sigma-Aldrich, Missouri, USA) as described (McLaughlin *et al*., 2023a). at the indicated time points (middle of day, end of day, end of night). Exudates were collected at 21 days after germination (dag) for 2 h, centrifuged to remove cellular debris, and stored at −80°C until analysis. Shoot and root fresh weight was recorded at the end of the experiment. Tissues were flash-frozen and stored at −80°C for metabolite analysis.

### Untargeted metabolite analysis

Root exudate volumes were normalized with 20mM Ammonium acetate according to total root mass. Tissue samples were ground at 1500 rpm for 1 min and extracted in 40:40:20 methanol: acetonitrile: ultrapure water twice for 1 h at 4°C with intermittent vortexing. Supernatants were dried under vacuum and dissolved at a concentration of 70 mg 200 µl^−1^ in 1:1 MeOH: ultrapure water. Samples were diluted 1:150 before metabolite analysis (McLaughlin *et al*., 2023b).

Two technical replicates were injected into a flow injection time of flight mass spectrometry system of an isocratic Agilent 1200 pump coupled to a Gerstel MPS2 autosampler and an Agilent 6550 QTOF mass spectrometer (Agilent, California, USA). MeOH and ammonium acetate samples were prepared as blanks, and a standard mixture of 1 µM amino acids was used as quality control (QC) samples injected at the beginning and end of each 96-well plate to ensure consistent peak heights throughout the run. After each sample group (QC, exudates, tissues), blank samples were injected to reduce the probability of sample carryover.

The platform operated with published settings (Fuhrer *et al*., 2011). The isocratic flow rate was 150 µl min^−1^ of mobile phase, consisting of 60:40% v/v isopropanol:water buffered with 5 mM ammonium fluoride for negative ionization mode. For online mass axis correction, homoserine and Hexakis (1H, 1H, 3H-tetrafluoropropoxy)-phosphazine were used. Mass spectra were recorded from 50-1000 m/z with a frequency of 1.4 spectra/s using the highest available resolving power. Source temperature was set to 325°C, with 5 ml/min drying gas and a nebulizer pressure of 30 spig. Fragmentor, skimmer, and octupole voltages were set to 175 V, 65 V, and 750 V, respectively. Spectral data was processed and annotated as published based on the Kyoto Encyclopedia of Genes and Genomes (KEGG) and Human Metabolome Database (HMDB) databases (Fuhrer *et al*., 2011).

Quality control samples were checked for m/z accuracy and intensity shifts. Data processing was completed in Visual code studio with Python. The two technical replicates for each sample were averaged (sample mean), and the mean of each species and negative control was calculated. Metabolites were deemed present in the dataset if the mean of the metabolite abundance in a single plant species was two times higher than the mean of negative control. The intensity of filtered metabolites underwent percentage normalization for PCA analysis. Statistical analysis (ANOVA and post hoc TUKEY tests) was completed for each metabolite present in the dataset. For the assignment of chemical classes to compounds, KEGG IDs were converted to InChIKeys, which was used as an input for Classyfire (https://cfb.fiehnlab.ucdavis.edu/, Djoumbou Feunang *et al*. (2016)). The following Python packages were used: pandas, numpy, scipy, matplotlib, sklearn, statsmodels, and multiprocessing.

### Sugar content measurement with HPAEC and sample collection

The growth medium was exchanged with 50 ml of ultrapure water for 2 h (similar to McLaughlin *et al*. (2023a)). Exudates were first collected at the end of the light/day period (EOD) at 19 dag and two days later at the end of the night period (EON). For shoot and root tissues, 50 mg of tissue fresh weight was collected at 21 dag MOD. Immediately after tissue collection, samples were incubated for 10 min at 80°C with 20 µl ethanol per mg of tissue. The supernatant was collected and dried under a vacuum at 23°C (UNIVAPO 100 ECH, UniEquip, Bayern, Germany). Samples were resuspended in 200 µl of deionized water.

Glucose, fructose, sucrose, raffinose, and trehalose in tissues and exudates were determined using the High-Performance Anion-Exchange chromatography with Pulsed Amperometric Detection (HPAECPAD, Thermo Fisher, Massachusetts, USA). 180 µL of all samples were added to sequential 1.5-mL columns of Dowex 50WX2 (cation exchange resin) and Dowex 1X8 (anion exchange resin) (203030025 (50WX2) and 203010025 (1X8), from Acros Organics by Thermo Fisher Scientific, Massachusetts, USA) to remove charged compounds, respectively. Uncharged metabolites, among them sugars, were eluted with 8 x 500 µl of MiliQ water, lyophilized, and redissolved in 110 µL of ultrapure water. 10 µl of a 1:100 dilution in ultrapure water was injected. Sugars were separated on a Dionex PA-20 column according to the following conditions: eluent A, water; eluent B, 100 mM NaOH; eluent C, 150 mM NaOH and 500 mM sodium acetate; eluent D, 800 mM NaOH. The gradient was as follows: 0 to 20 min, a rising gradient of 25% B to 50% B (respectively 75% A and 50% A) was established; 20 to 30 min, 50% B and 50% A; 30 to 31 min, 100% C; 31 to 41 min, 100% C; 41 to 64 min, 25% B and 75% A. Peaks were identified by coelution with glucose, fructose, sucrose, raffinose, and trehalose chemical standards. Peak areas were determined using the Chromeleon software. Methods from Fulton *et al*. (2008).

### Total organic carbon quantification

Root exudate samples were collected individually for each seedling in 20 mM ammonium acetate (102308585, Sigma-Aldrich, Missouri, USA) for 2 h and analyzed with an optical density plate reader (plate reader Biotek Epoch2, Agilent Technologies, California, USA) modified from a published method (Oburger *et al*., 2022) developed for quantification of low carbon levels in root exudates. Luminescence was determined using the BioTek Gen 5 software (Agilent Technologies, California, USA). Briefly, a standard curve was made with potassium hydrogen phthalate (KHP, 1003326003, Sigma-Aldrich, Missouri, USA) and ammonium acetate (102308585, Sigma-Aldrich, Missouri, USA), and absorbance was determined at 260 nm. Exudate samples were quantified undiluted. Data was visualized as carbon per root fresh weight.

### ROS production assay

To measure ROS production, *Arabidopsis thaliana* Col-0 seedlings were grown as described above for 9, 14, and 21 days on ½ MS plates with and without 1% sucrose in sterile conditions. For 9 d and 14 d-old plants, the whole seedling was dipped into deionized water to remove the MS excess and transferred to 96-well plates. For the older 21 d old plant, leaf discs of 0.4 mm in diameter were punctured with a biopsy punch and moved to the 96-well plate. 150 µl deionized water was added to all 96-well plates, followed by incubation in the dark for 24 h. The next day, the water was replaced with a solution containing 10 µl ml^−1^ horseradish peroxidase (HRP), 100 nM flg22 (SciLight Biotechnology LLC), and 0.5 µM L-012 for seedlings and 0.5 µM luminol for leaf discs. Chemiluminescence of the ROS burst was visualized under a Tecan plate reader (Spark Microplate reader, Tecan, Männedorf, Germany) for 30 min. Chemiluminescence was determined using the Tecan Sparkcontrol software. Seedlings were weighed immediately after the assay and their fresh weight was used to normalize the ROS curve for 9 d and 14 d-old ROS bursts.

### RNA extraction and sequencing

*Arabidopsis thaliana* Col-0 seedlings were grown on plates supplemented with and without sucrose 1% for 9 d, 14 d, and 21 d. RNA was extracted from whole seedlings (RNeasy Plant Mini kit, Qiagen, USA). The Functional Genomics Center Zurich prepared the RNAseq libraries and performed the sequencing: The quality of the isolated RNA was determined with a Qubit® (1.0) Fluorometer (Life Technologies, California, USA) and a Fragment Analyzer (Agilent, Santa Clara, California, USA). Only those samples with a 260 nm/280 nm ratio between 1.82.1 and a 28S/18S ratio within 1.52 were further processed. The TruSeq Stranded mRNA (Illumina, Inc, California, USA) was used in the succeeding steps. Briefly, total RNA samples (100-1000 ng) were polyA enriched and then reverse-transcribed into double-stranded cDNA. The cDNA samples were fragmented, end-repaired, and adenylated before the ligation of TruSeq adapters containing unique dual indices (UDI) for multiplexing. Fragments containing TruSeq adapters on both ends were selectively enriched with PCR. The quality and quantity of the enriched libraries were validated using Qubit® (1.0) Fluorometer and the Fragment Analyzer (Agilent, Santa Clara, California, USA). The product is a smear with an average fragment size of approximately 260 bp. The libraries were normalized to 10nM in Tris-Cl 10 mM, pH 8.5 with 0.1% v/v Tween20. The NovaseqX (Illumina, Inc, California, USA) was used for cluster generation and sequencing according to standard protocol. Sequencing was paired end at 2 x150 bp.

Analysis of the read data was performed using the SUSHI framework (Hatakeyama *et al*., 2016). Reads were preprocessed with fastp to trim adapters (Chen *et al*., 2018). Read alignment was done using the STAR aligner (Dobin *et al*., 2013) with the TAIR10 reference. Gene-level expression values were generated using the featureCounts function of the R package Rsubread (Liao *et al*., 2013).

### RNAseq and GO term analysis

Reads Per Kilobase Million (RPKM) data were used for a principal component analysis. The non-normalized sequence read counts were processed with the Python DESeq2 workflow (https://github.com/owkin/PyDESeq2), a Python implementation of the DESeq2 pipeline for bulk RNAseq for differential expression analysis (Love *et al*., 2014; Zhu *et al*., 2019).

The heatmap is based on the raw counts data of 9 d, 14 d, and 21 d samples. The heatmap was generated using the seaborn library (sns.clustermap). The clustering was performed using the Bray-Curtis dissimilarity metric and the average linkage method. A log base 10 transformation was applied to stabilize and normalize the distribution. This transformation was performed on the entire dataset with an offset of 1 added to avoid undefined values for zero entries.

The Metascape GO-term classification system (Zhou *et al*., 2019) and the PANTHER GO-slim classification system (Mi *et al*., 2019) were used to attribute GO terms.

### Infection assay with *Pseudomonas syringae* DC3000

Plants were grown on soil for 3 weeks at 18°C day 8 h/ 22°C 16 h night with 150-160 mmol m^−2^ s^−1^ illumination. *Pseudomonas syringae* DC3000 (COR-) was cultivated on Kings B medium supplemented with 50 µg ml^−1^ rifampicin for 24 h at 28°C. One day before the infection assay, scraped bacteria from the plate were cultured overnight in a liquid culture supplemented with 50 µg ml^−1^ rifampicin at 28°C with shaking. The next day, bacterial cells were collected by centrifugation at 4000 g for 5 min and resuspended in 10 mM MgCl_2_ until reaching an optical density at 600 nm of 0.2. The final solution was sprayed on and below the rosette leaves. After three days of incubation, 3 leaf discs of 0.4 mm diameter per plant were collected with a biopsy puncher. They were ground with a Genogrinder for 1.5 min at 1500 rpm with 200 µl 10 mM MgCl_2_. The solution was then serially diluted and 20 µl of each dilution series was dispensed on a Luria-Bertani (LB) agar plate supplemented with 50 µg ml^−1^ rifampicin. Plates were incubated overnight at 28°C and colony counting was performed the next day.

### Bacterial growth in root exudate media

50 ml exudates collected at the end of the day and the end of the night for the sugar concentration determination of *Arabidopsis thaliana* Col-0 and mutants were lyophilized and resuspended with ultrapure water to concentrate 150x. Exudates were filter sterilized with a 0.22 µm filter (PALL Life Sciences, USA). *Escherichia coli*, *Pseudomonas simiae*, *Paraburkholderia phytofirmans*, *Bacillus subtilis*, and *Pseudomonas syringae* were cultured in tryptic soy broth (TSB) at 28°C. Bacteria were collected by centrifugation at 4000 g for 5 min and resuspended in ½ MS to an optical density of 0.4 at 600 nm. 96-well plates were filled with 50 µl of 150x concentrated exudates and 50 µl of bacterial suspension. Optical density was determined at 600 nm each hour for 36 h. As a positive growth control, TSB medium was added to the plates. As a negative growth control, bacterial suspensions were mixed with ½ MS, or with experimental blanks (growth jars without plants). If the area under the curve (AUC) of bacterial growth in negative samples was significantly lower than the growth on root exudates samples, bacteria were considered to grow (based on T-Test vs. negative samples).

To confirm that bacterial growth was not only supported by ½ MS (the dilution solution for bacteria), the AUC of bacteria grown on ½ MS was subtracted from the AUC of the corresponding bacteria grown on root exudates.

To compare the bacterial growth between Col-0 and mutants, the AUC of bacteria grown on Col-0 was subtracted from the AUC of the corresponding bacteria grown on each mutant. Each growth setup was performed in triplicates.

## Supplementary material

**Figure S.1:**
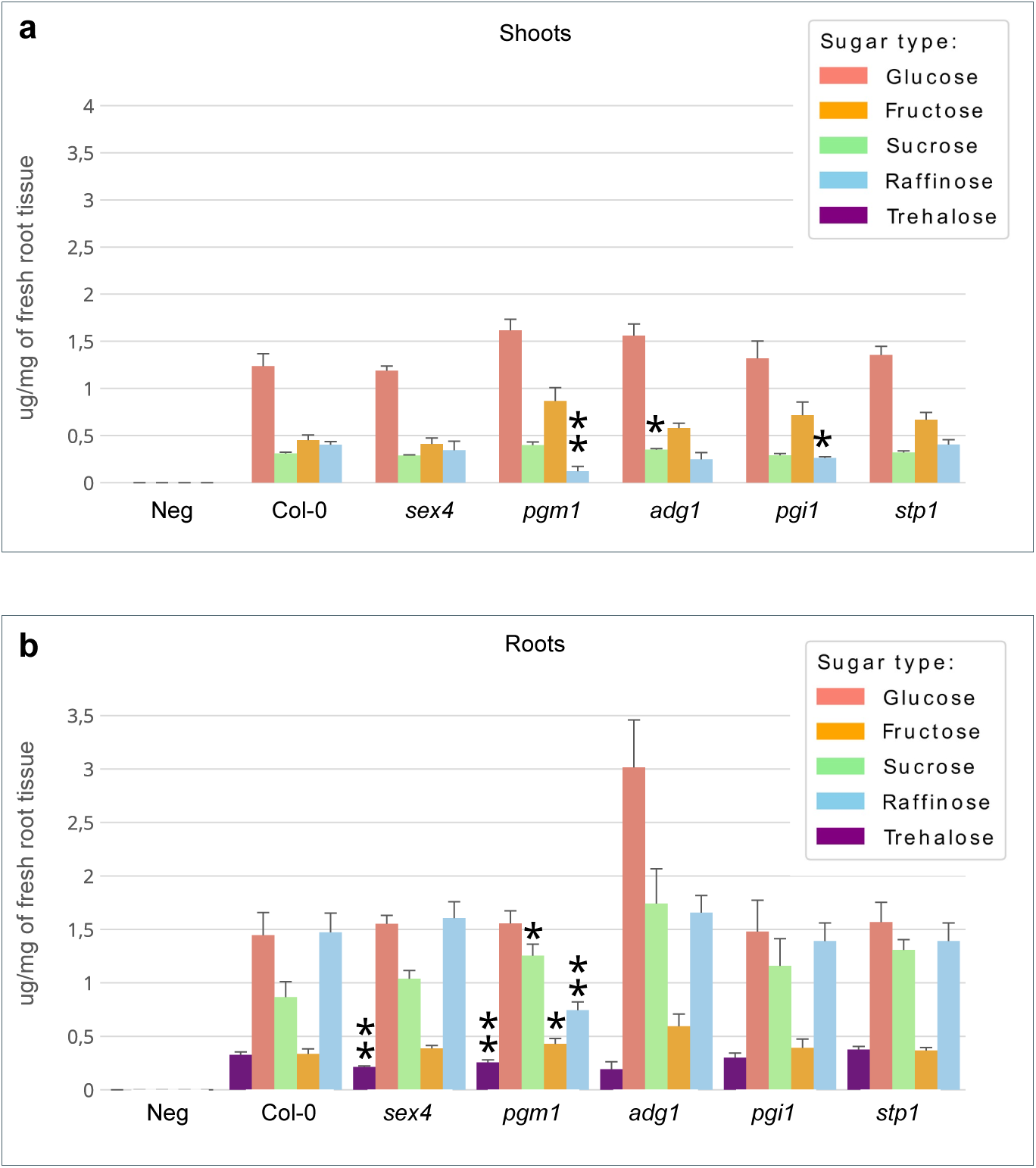
Sugar content in tissues. (a-b) Sugar content of shoots (a) and roots (b) of sex4, pgm1, adg1, pgi1, and stp1. Data is displayed as average *±* S.E.M., *: pvalue < 0.05, **: pvalue < 0.01 (T-Test vs. Col-0 corresponding sugar). Colors for different sugars: pink: glucose, orange: fructose, green: sucrose, blue: raffinose, purple: trehalose.

**Figure S.2:**
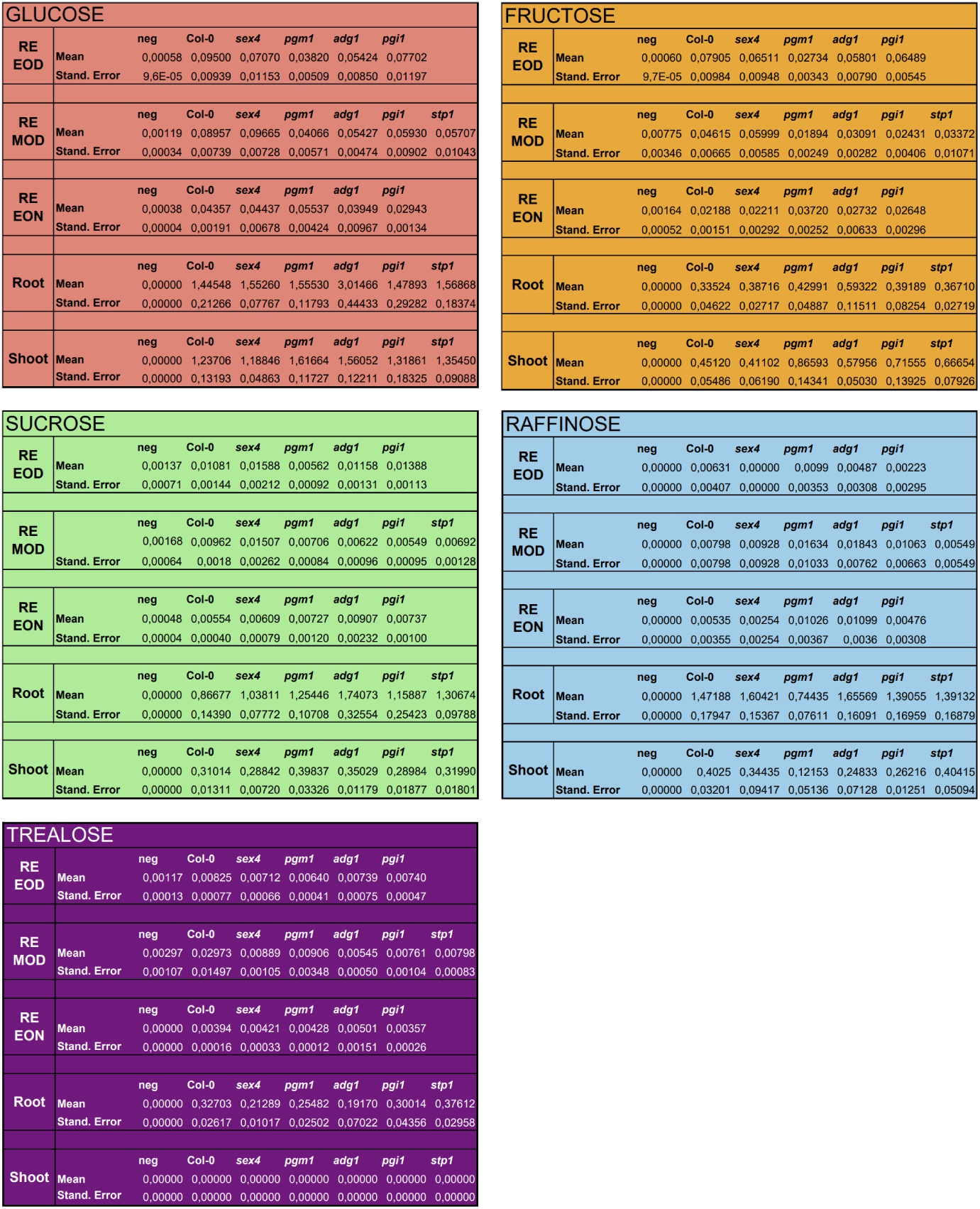
Sugar content in tissues and exudates. Raw data of sugar content in tissues and exudates. Color code: pink: glucose, orange: fructose, light green: sucrose, blue: raffinose, purple: trehalose. RE EOD: root exudates at the end of day, RE MOD: root exudates at midday, RE EON: root exudates at the end of night. The mean values come are based on 5 biological replicates (jars with each 5 Arabidopsis individuals). Stand. Error: standard error.

**Figure S.3:**
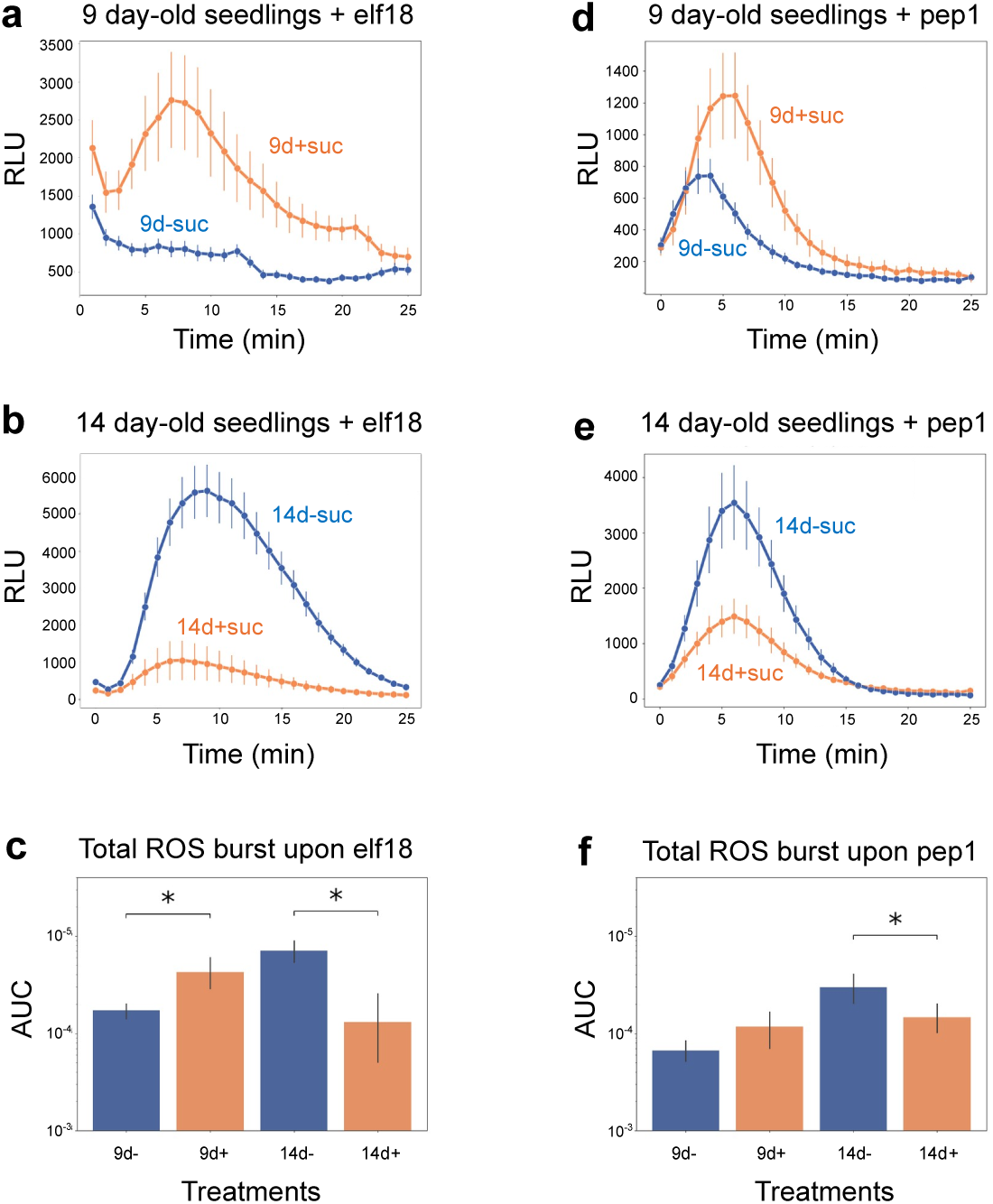
Reactive oxygen species production upon elf18 and pep1 elicitors treatment. (a-b) Reactive oxygen species (ROS) production for 9 d (a), 14 d (b) upon elf18 (100 nM) treatment. (d-e) Reactive oxygen species (ROS) production for 9 d (a), 14 d (b) upon pep1 (500 nM) treatment. (c and f) Area under the curve for all samples in a and b and d and e respectively. Data is based on 1 biological replicate. Data is displayed as average *±* S.E.M., *: pvalue < 0.05 (T-Test with vs without sucrose samples). Plants were grown on ½ MS (Murashige and Skoog) plates with or without supplementation of 1% sucrose (orange and blue curves, respectively). +: with sucrose, −: without sucrose. RLU: relative light units. AUC: area under the curve.

**Figure S.4:**
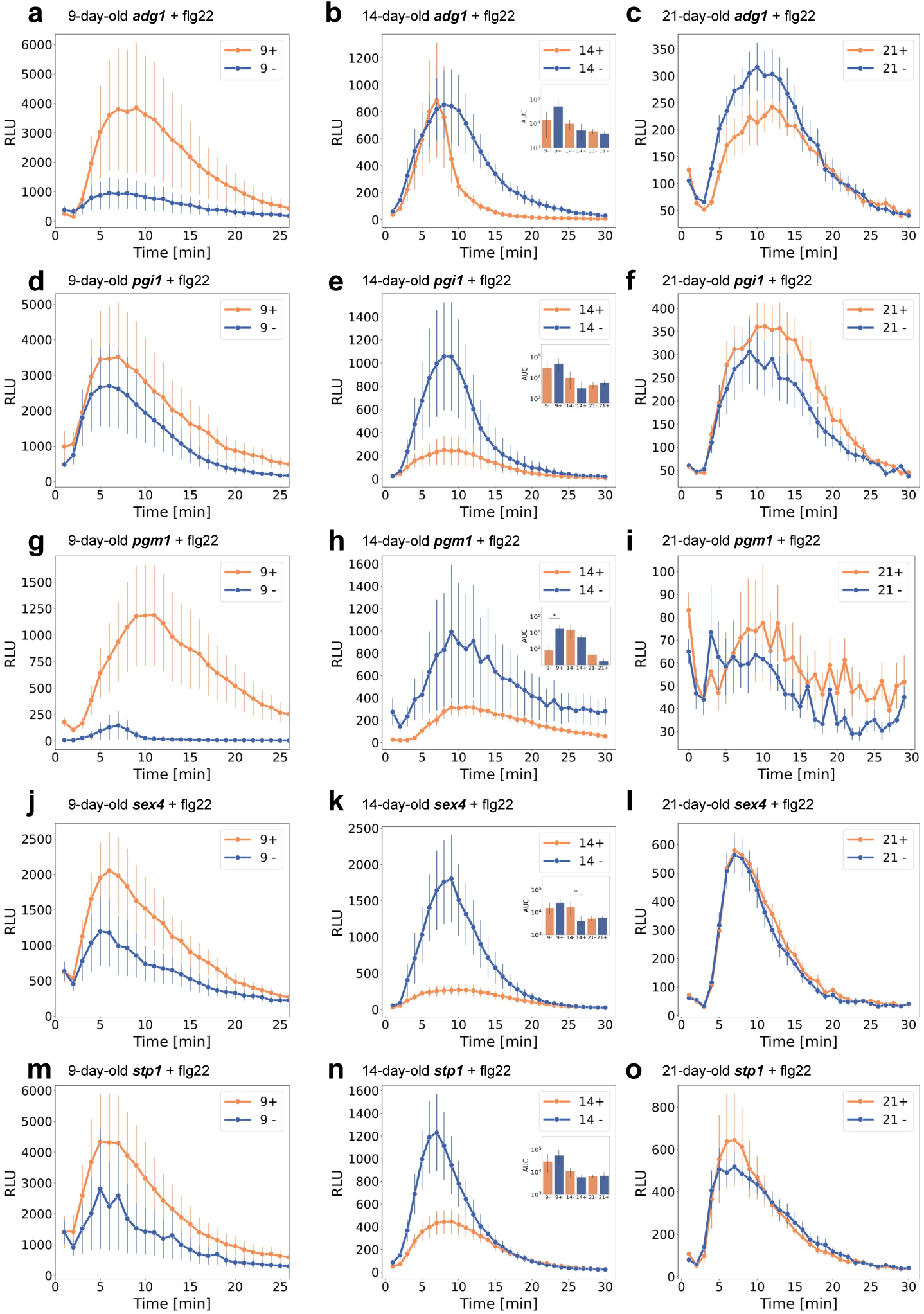
Reactive oxygen species production upon flg22 elicitors treatment on sugar metabolism mutant lines. (a-o) Reactive oxygen species (ROS) production of sugar metabolism mutant lines. Plants were grown on ½ MS (Murashige and Skoog) plates with or without supplementation of 1% sucrose (orange and blue curves, respectively) and treated with flg22 (100 nM). (a-c) *adg1* at 9 d (a), 14 d (b) and 21 d (c). (d-f) *pgi1* at 9 d (d), 14 d (e) and 21 d (f). (g-i) *pgm1* at 9 d (g), 14 d (h) and 21 d (i). (j-l) *sex4* at 9 d (j), 14 d (k) and 21 d (l). (m-o) *stp1* at 9 d (m), 14 d (n) and 21 d (o). AUC of the respective 3 time points are displayed in reduced size in 14 d treatment for each mutant (b, e, h, k and n). Data is based on 1-2 biological replicates. Data is displayed as average *±* S.E.M., *: pvalue < 0.05 (T-Test with vs without sucrose samples). +: with sucrose, −: without sucrose. RLU: relative light units. AUC: area under the curve.

**Figure S.5:**
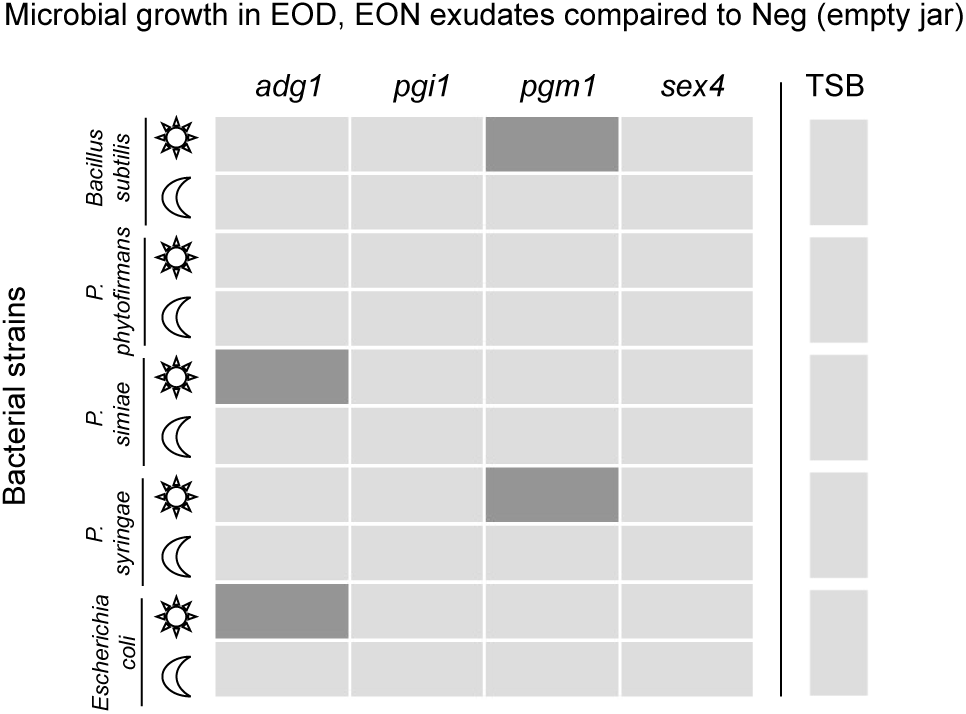
Neg comparison heatmap for microbial growth assay. Binary heatmap of statistically significant pvalue of the area under the curve of bacteria growing on mutant exudates vs. Neg exudates (non-plant jars). Color code: light gray: pvalue < 0.05 (bacteria growth on mutant exudates is different from bacteria growth on Neg exudates), dark gray: pvalue 0.05 (bacteria growth on mutant exudates is not different from bacteria growth on Neg exudates). pvalue < 0.05 (T-Test mutant exudates vs Neg exudates). Microbes: *Bacillus subtilis*, *Paraburkholderia phytofirmans*, *Pseudomonas simiae*, *Pseudomonas syringae* DC3000, *Escherichia coli*. Tryptic soy broth (TSB) was used to confirm bacterial activity.

